# Empirical estimation of marine phytoplankton assemblages in coastal and offshore areas using an *in situ* multi-wavelength excitation fluorometer

**DOI:** 10.1101/2021.08.31.458334

**Authors:** Taketoshi Kodama, Yukiko Taniuchi, Hiromi Kasai, Tamaha Yamaguchi, Misato Nakae, Yutaka Okumura

**Author notes:** Corresponding author (TK).

## Abstract

Phytoplankton assemblages are important for understanding the quality of primary production in marine ecosystems. Here, we describe development of a methodology for monitoring marine phytoplankton assemblages using an *in situ* multi-wavelength excitation fluorometer (MEX) and its application for seasonal observations in coastal and offshore areas around Japan. The MEX recorded the fluorescence excited with nine light-emitting diodes, temperature, and sensor depth. We prepared reference datasets comprising temperature, MEX fluorescence, and plant-pigment-based chemotaxonomy phytoplankton assemblages. Target MEX fluorescence was decomposed by reference MEX fluorescence using a linear inverse model for calculating coefficients after the reference data were limited by temperature, followed by reconstruction of plant-pigment-based chemotaxonomy of the target MEX fluorescence using the coefficients and the chemotaxonomy assemblages of the reference data. Sensitivity analysis indicated poor estimation of the proportion and/or chlorophyll *a*-based abundance of chlorophytes, haptophytes, prasinophytes, and prochlorophytes; however, limiting the estimations to five chemotaxonomic groups [diatoms, dinoflagellates, cryptophytes, cyanobacteria (cyanophytes and prochlorophytes), and other eukaryotes (chlorophytes, haptophytes, and prasinophytes)] resulted in positive correlations of both the proportion and abundances, suggesting that the five taxonomic abundances were well-estimated using the MEX. Additionally, MEX observations denoted spatial and seasonal variations of phytoplankton assemblages, with high contributions from other eukaryotes in every area and season, cyanobacteria highly during the summer in surface Kuroshio and Japan Sea waters, and diatoms in the Oyashio and Oyashio–Kuroshio transition areas and the Okhotsk Sea. Furthermore, ratios of water-column-integrated chlorophyll-based abundances to those on the surface at the chemotaxonomy group level were differed among the areas and groups. These findings suggested that phytoplankton-assemblage monitoring in the vertical direction is essential for evaluation of their current biomass, and that the MEX promotes the acquisition of these observations.

## Introduction

Phytoplankton abundance and composition are important parameters for understanding marine ecology [1], with differences in phytoplankton assemblages linked to differences in higher-trophic marine-organism assemblages [2]. Additionally, phytoplankton assemblages affect biogeochemical element cycles. For example, diatoms change dissolved Si into particulate Si, and the carbon-isotope fraction of tuna (top predators in the ocean) in these the previous two decades suggest that the phytoplankton community has changed from ^13^C-rich to ^13^C-poor at the global scale [3]. Lorrain et al. [3] suggested that diatoms are decreasing in oceans; however, evidence regarding changes in the phytoplankton community is limited. Therefore, monitoring of phytoplankton assemblages is essential at both the global and local scales, and consistent analysis methods are required [4].

Microscopic observation is the most basic method for monitoring phytoplankton assemblages. Chemotaxomic identification using photosynthesis and photoprotective marker pigments represents an alternative and more convenient method [1]; however, chemical analyses during plant-pigment identification using high-performance liquid chromatography (HPLC) is neither safe nor environmentally friendly based on the use of organic solvents for pigment extraction. Moreover, HPLC measurement requires ≥30 min, and plant-pigment markers degrade easily [5], which prevents routine and high-resolution observations. By contrast, ultra-HPLC required a short run time (7 min) [6], although high-resolution observations are difficult when considering on-board operations.

*In situ* multi-excitation chlorophyll fluorimetry (using a submersible spectrofluorometer) is a promising technique for rapidly evaluating phytoplankton-community structure [7]. Fluoroprobe (bbe Moldaenke, Schwentinental, Germany) is a dominant submersible spectrofluorometer for monitoring phytoplankton communities in the ocean [8]. Fluoroprobe uses five light-emitting diodes (LEDs; operating at 470, 525, 570, 590, and 610 nm) to excite accessory pigments associated with the photosystem II antenna system [8] and allows detection of the chlorophyll-based abundance of four groups (green, blue, brown, and mixed) [7]. The Multi-Exciter (MEX; JFE-Advantech, Hyogo, Japan) is an alternate submersible spectrofluorometer that uses nine LEDs (375, 400, 420, 435, 470, 505, 525, 570, and 590 nm) to excite the accessory pigments associated with the photosystem II antenna system and can estimate the chlorophyll-based abundance of three groups (green algae, brown algae, and cyanobacteria) as the default [9]. However, Wang et al. [10] indicated that the default conversion method from the fluorescence to the phytoplankton assemblages using the MEX software is imprecise at the study site, and that site-specific methods are required for better estimations. Although the conversion techniques described by Wang et al. [10] were adequate for estimating phytoplankton assemblages from fluorescence measurements in the East China Sea, they did not describe the specific details of those techniques. Therefore, MEX observations are not usually converted to phytoplankton-community structures, as described by Fujiwara et al. [11].

In this study, we developed methods to rapidly evaluate high-quality phytoplankton assemblages using an open-source computer program with the MEX. Specifically, we used an empirical method to convert MEX data to phytoplankton compositions and evaluated this conversion method using sensitivity analysis. To apply this method, we prepared reference data from the observations and calculated spatiotemporal variations of 1-m pitch vertical profiles of phytoplankton-community structure in the coastal and offshore areas of the ocean around Japan. Using this data, we then compared the surface abundances and water-column integrated abundances in order to determine the representativeness of the surface environments in the water column, which can be observed with satellites [4].

## Materials and Methods

### Field MEX observations

The MEX observations were conducted at 366 stations during 23 cruises around Japan in 2016, 2019, and 2020 (Fig 1). The areas were in the Japanese closed sea, Japanese EEZ, and high seas; we need not require any permission for observation in these areas except Japanese government, and our cruises were permitted Japan Fisheries Research and Education Agency, and Fisheries Agency of Japan. The observation areas were divided into four areas based on their geographic characteristics: the Kuroshio and Oyashio areas (including the Oyashio–Kuroshio transition area), the Okhotsk Sea (Okhotsk), and the Sea of Japan (JS). The MEX observations in 2016 were only conducted in the JS at 29 stations, and in 2019 and 2020, they were conducted in four areas. When winter, spring, summer, and autumn were defined as occurring from December to February, March to May, June to September, and October to November, respectively, observations in the Kuroshio area were conducted throughout the four seasons, and observations were not conducted in autumn and summer in the JS or Oyashio areas, respectively. In Okhotsk, observations were only conducted in June (including two stations conducting observations on 31 May 2019) and September in both 2019 and 2020; thus, the results of June were categorized as from the spring. The seasonal and spatial numbers of the stations are listed in Table 1.

**Fig 1.**
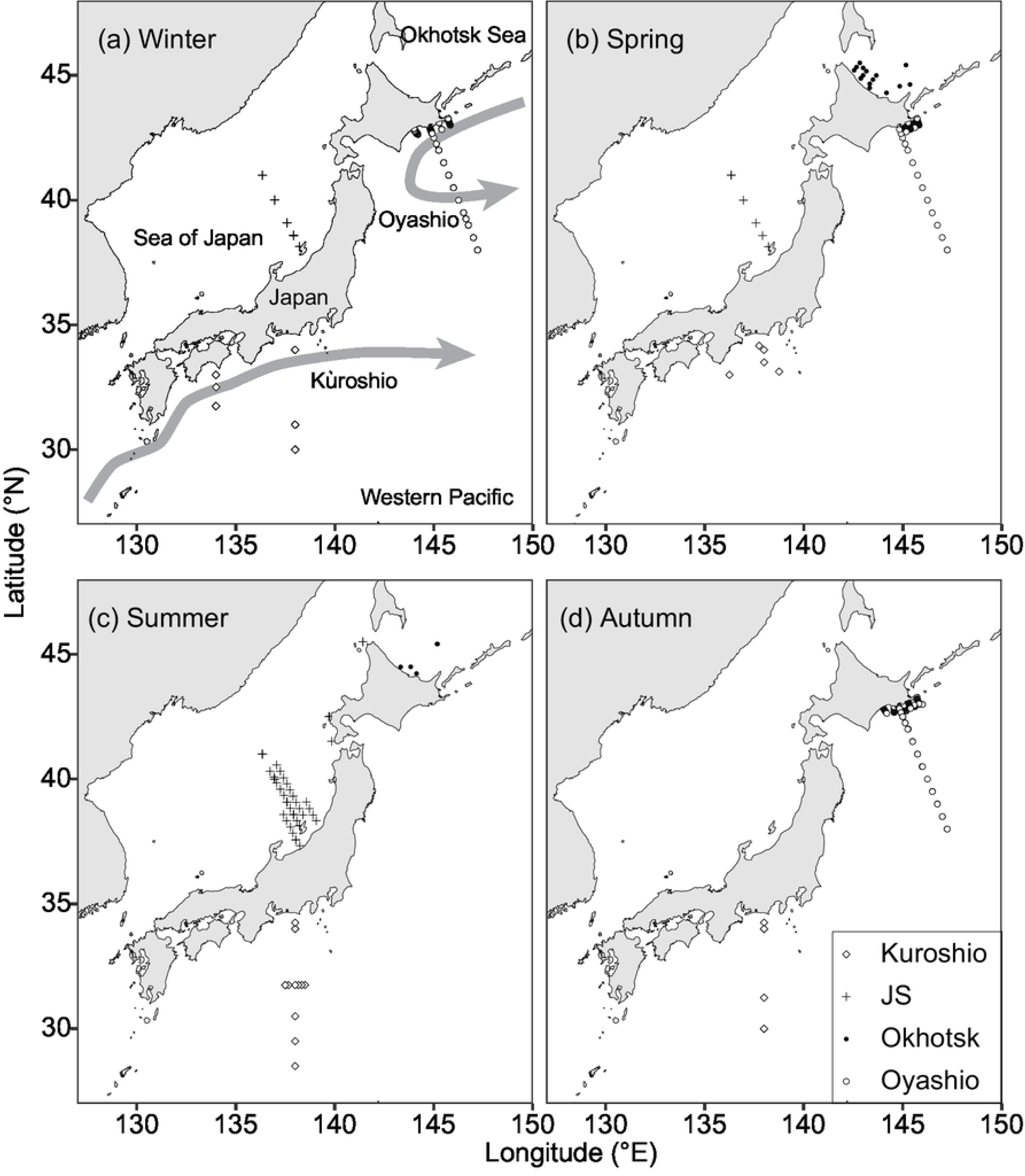
Maps of MEX (Multi-Exciter) observations. Maps show observations in (a) winter, (b) spring, (c) summer, and (d) autumn. Different symbols denote different areas: diamond, Kuroshio; cross, Sea of Japan (JS); closed circle, Okhotsk Sea; and open circle, Oyashio, including the Oyashio–Kuroshio transition. Schematic flows of Kuroshio and Oyashio are denoted as arrows.

**Table 1.** List of Numbers of Stations Sampling MEX (Multi-Exciter) Observations and the Number of Samples Used for Discrete Plant-pigment Analysis.

|  | Winter | Spring | Summer | Autumn |
| --- | --- | --- | --- | --- |
| MEX (Multi-Exciter) |  |  |  |  |
| Kuroshio | 6 | 5 | 13 | 6 |
| JS (Sea of Japan) | 5 | 7 | 45 | 0 |
| Okhotsk | 0 | 16 | 5 | 0 |
| Oyashio | 62 | 77 | 0 | 121 |
| Discrete plant pigments |  |  |  |  |
| Kuroshio | 27 | 18 | 45 | 13 |
| JS | 16 | 26 | 48 | 0 |
| Okhotsk | 0 | 14 | 1 | 0 |
| Oyashio | 17 | 13 | 0 | 33 |

Observations of the Okhotsk area conducted in June were considered as spring observations.

These observations were conducted using three MEXs. The MEX excited the matter in the water using nine LED light sources (375, 400, 420, 435, 470, 505, 525, 570, and 590 nm), and the binary data recorded in the MEX were transformed into numerical values using the default software attached to the MEX. The MEXs were calibrated at the factory before shipping, and we evaluated difference between the three MEXs at every excitation source by using 10 types of algal culture collections (diatoms, cryptophytes, chlorophytes, and *Synechococcus*) in December 2019.

Fluorescence measurements were obtained by the MEX recording the temperature and pressure (depth of the sensor), with MEX recordings not as sensitive as those of SBE9plus or SBE19 conductivity, temperature, and depth sensors (Sea-Bird Scientific, Bellevue, WA, USA) but sufficient for subsequent analyses. Although capable of recording these values every 0.1 s, for the present study, values were recorded every 1 s, with those recorded every 0.1 s averaged with those recorded every 1 s. Data from a sensor depth of ≥2 m over 300 s were categorized as “observations”, because the sensor sometimes recorded a depth of 1 m on the deck. This allowed the MEX observations to be consistent with field-book recorded observations; therefore, the MEX-based results at the surface indicated those at 2-m depth. We converted the 1-s MEX data to 1-m pitch data using the pressure data of the MEX.

To estimate the water-column-integrated values, the 1-m pitch data were integrated from depths of 2 m to 100 m, except in the Kuroshio, where maximal subsurface chlorophyll was observed below a depth of 100 m. Thus, data from depths of 2 m to 200 m were integrated as water-column-integrated values.

### Plant-pigment-based plankton abundance

Samples for plant-pigment analysis were collected in 2019 and 2020 at stations using MEX for observation. Particles in 0.5 L to 2.3 L of seawater were collected on 0.7-µm mesh glass-fiber filters (Whatman GF/F) under gentle suction (<0.02 MPa). The filter was immersed in 1 mL of *N,N*-dimethylformamide (DMF) and stored in the dark at <−20°C until onshore analysis or rapidly frozen in liquid nitrogen without DMF immersion and stored at <70°C until onshore analysis. The former storage method can maintain the status of chlorophyll and accessory pigments (except diadinoxanthin) for 180 days [12], and the latter method is the conventional method for plant-pigment analysis. We collected 271 samples from four areas at depth of between 10 m and 170 m (Table 1).

The extracted plant-pigment concentrations were measured using an HPLC system (Shimadzu, Kyoto, Japan). DMF (1 mL) was added to samples stored without DMF, and after filter removal, the extracts were centrifuged at a maximum speed of 17,000*g* for 10 min, followed by injection of 300 µL of supernatant into the HPLC system after mixing with 90 µL ultrapure water. Plant pigments were analyzed according to the protocol described by Zapata et al. [13]. During analysis, a 4.6 mm × 150-mm reversed-phase C_8_ column (Symmetry 3.5 μm; Waters, Milford, MA, USA) with a guard column (Symmetry Sentry; Waters) was maintained at 25°C. Plant pigments were detected using a photodiode array UV-Vis detector (SPD-M10AV; Shimadzu). Twenty types of major plant pigments were identified by their retention times and absorption spectra and quantified by the peak areas according to reference standards (Danish Hydraulic Institute, Hørsholm, Denmark). The following 13 types of plant pigments were detected using the following analyses: peridinin, 19’- butanoyloxyfucoxanthin, fucoxanthin, 19’-hexanoyloxyfucoxanthin, neoxanthin, prasinoxanthin, violaxanthin, alloxanthin, lutein, zeaxanthin, chlorophyll *b*, monovinyl chlorophyll *a* (MVChla), and divinyl chlorophyll *a* (DVChla). The sum of MVChla and DVChla concentrations was described as total chlorophyll *a* concentration (TChla).

Plant-pigment-based chemotaxonomic phytoplankton assemblages were calculated using a Bayesian compositional linear inverse technique [14] using R software [15]. For the calculation, we used the “limSolve” package [16], which is comparable to the Mackey et al. [1] program [14]. Some plant-pigment ratios were previously published for the calculation of chemotaxonomic phytoplankton compositions using the CHEMTAX program in our study area [17, 18]; however, the plant-pigment ratios of eight algal classes [prasinophytes, dinoflagellates, cryptophytes, haptophytes (types 3 and 4), chlorophytes, cyanobacteria, and diatoms] were calculated using the default settings for the “limSolve” package [1], and those of prochlorophytes were calculated as described by Nishibe et al. [19]] (Table 2).

**Table 2.** Ratios of Biomarker Pigments According to the Linear Inverse Technique.

|  | Per | But | Fuc | Hex | Neo | Pra | Vio | All | Lut | Zea | Chlb | MVChla | DVChla |
| --- | --- | --- | --- | --- | --- | --- | --- | --- | --- | --- | --- | --- | --- |
| Prasinophytes | 0 | 0 | 0 | 0 | 93.0 | 113.1 | 104.7 | 0 | 87.7 | 0 | 96.8 | 99.8 | 0 |
| Dinoflagellates | 114 | 0 | 0 | 0 | 0 | 0 | 0 | 0 | 0 | 0 | 0 | 87.9 | 0 |
| Cryptophytes | 0 | 0 | 0 | 0 | 0 | 0 | 0 | 100.6 | 0 | 0 | 0 | 99.9 | 0 |
| Haptophytes 3 | 0 | 0 | 0 | 109.7 | 0 | 0 | 0 | 0 | 0 | 0 | 0 | 83.6 | 0 |
| Haptophytes 4 | 0 | 91.6 | 94.4 | 97.6 | 0 | 0 | 0 | 0 | 0 | 0 | 0 | 106.7 | 0 |
| Chlorophytes | 0 | 0 | 0 | 0 | 113.5 | 0 | 19.4 | 0 | 119.8 | 111.1 | 105.4 | 93.1 | 0 |
| Diatoms | 0 | 0 | 97.2 | 0 | 0 | 0 | 0 | 0 | 0 | 0 | 0 | 102.1 | 0 |
| Cyanophytes | 0 | 0 | 0 | 0 | 0 | 0 | 0 | 0 | 0 | 118.6 | 0 | 93.5 | 0 |
| Prochlorophytes | 0 | 0 | 0 | 0 | 0 | 0 | 0 | 0 | 0 | 32.0 | 111 | 0 | 100 |

Per: peridinin; 19'-butanoyloxyfucoxanthin; Fuc: fucoxanthin; Hex: 19'-hexanoyloxyfucoxanthin; Neo: neoxanthin, Pra: prasinoxanthin;
Vio: violaxanthin; All: alloxanthin, Lut: lutein, Zea: zeaxanthin, Chlb: chlorophyll *b*; MVChla: monovinyl chlorophyll *a*; and DVChla: divinyl chlorophyll *a*.

We evaluated the effects of the fluorescence values of nine LEDs and temperature on plant-pigment-based phytoplankton chemotaxonomy using permutational multivariate analysis of variance (PERMANONA) in the “vegan” package [20] in R. The distances between samples were calculated based on the Bray– Curtis method, and the number of permutations was 999. The effects of the fluorescence values and temperature were checked using the determination coefficient (*R*^2^).

The TChla was also estimated from MEX fluorescence using fluorescence and sensor depth. The intercept of multiple regression analysis was set to zero in order to avoid the negative predicted TChla. The Pearson’s matrix indicated that TChla (MVChla + DVChla) was most strongly correlated with fluorescence excited at 400 nm (*r* = 0.779), followed by that at 435 nm (*r* = 0.778). The Akaike information criterion (AIC) indicated that the following equation was optimal for predicting TChla from MEX fluorescence (depth was not selected as an explanatory variable based on the AIC selection):

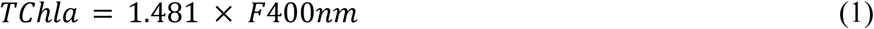

where *F_400nm_* represents fluorescence excited at 400 nm.

### Conversion of MEX data to phytoplankton assemblage

Conversion from MEX fluorescence values to chemotaxonomic phytoplankton assemblages was performed using the linear inverse technique and R software [15]. Additionally, calculating chemotaxonomic phytoplankton assemblages from pigment data was performed using the package “limSolve” [16]. We did not apply the Bayesian compositional linear inverse technique in order to save calculation resources.

Conversion of the observed MEX data (*f*) to the proportions of phytoplankton assemblages (*p*) was calculated in three steps. Before conversion, we prepared a reference database (*U*) comprising MEX fluorescence (*F*_1_ –*F*_271_), temperature (*T*_1_ –*T*_271_), and plant-pigment-based chemotaxonomic phytoplankton assemblage (*P*_1_ –*P*_271_) data collected from the same water. The MEX fluorescence was standardized to a summary of nine light-source emission fluorescence values of 1. We obtained sufficient reference data (*n* = 271) and created the subset of reference data (*A*) using *f*_T_, as follows:

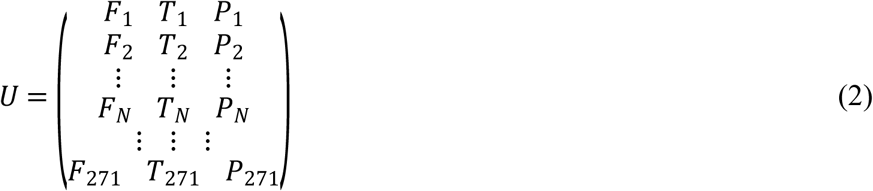

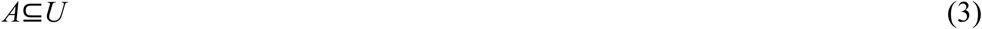

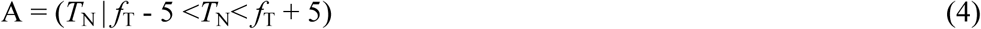

where N represents a natural number, except 0.

The observed MEX fluorescence for conversion (*f*) was then decomposed using the reference MEX fluorescence (*F*_1_ –*F*_271_):

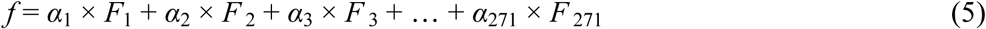

where the coefficient α_n_ was a number between 0 and 1, and the coefficients were established such that the residual was the lowest. After limits were established for Eqs. (2) through (4), the coefficient for Eq. (5) (*α*_n_) was 0 when the temperature of the reference data (*T*_N_) was within the range of ±5°C for the transformed values (*f*_T_ or *f*_D_), respectively. We then calculated *p* (phytoplankton assemblage) using the coefficients (*α*_n_), as follows:

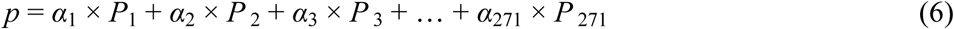

After these steps, *f* was converted to *p,* and the open-source R script and the reference dataset (*U*) were used in the data repository.

We considered that the sensor depth was used to limit the reference data (i.e., for the creation of *A*); however, we observed serious gaps in the vertical profiles of the MEX-estimated phytoplankton assemblages, with these gaps occurring infrequently when the reference data were not limited to depth. Therefore, the sensor depth was not included in Eq. (4).

## Results

### Sensitivity analyses

We conducted sensitivity analyses to evaluate the robustness of the conversion process in Eqs. (5) and (6). When conversion was conducted based on reference data from the same stations, the proportion of each chemotaxomic group based on MEX data was largely the same as that of the reference data (Fig 2). Therefore, the conversions were considered heavily dependent on the reference data collected at the same site. Therefore, in the sensitivity analysis, *F*_N_ was decomposed using reference database *F* but not from data collected at the same station. The estimated phytoplankton assemblage (*p*) was then compared to observational values (*P*_N_). When we observed a significant correlation between *p* and *P*, the conversion processes in Eqs. (5) and (6) were considered robust. Additionally, we compared the proportion and chlorophyll-based abundance.

**Fig 2.**
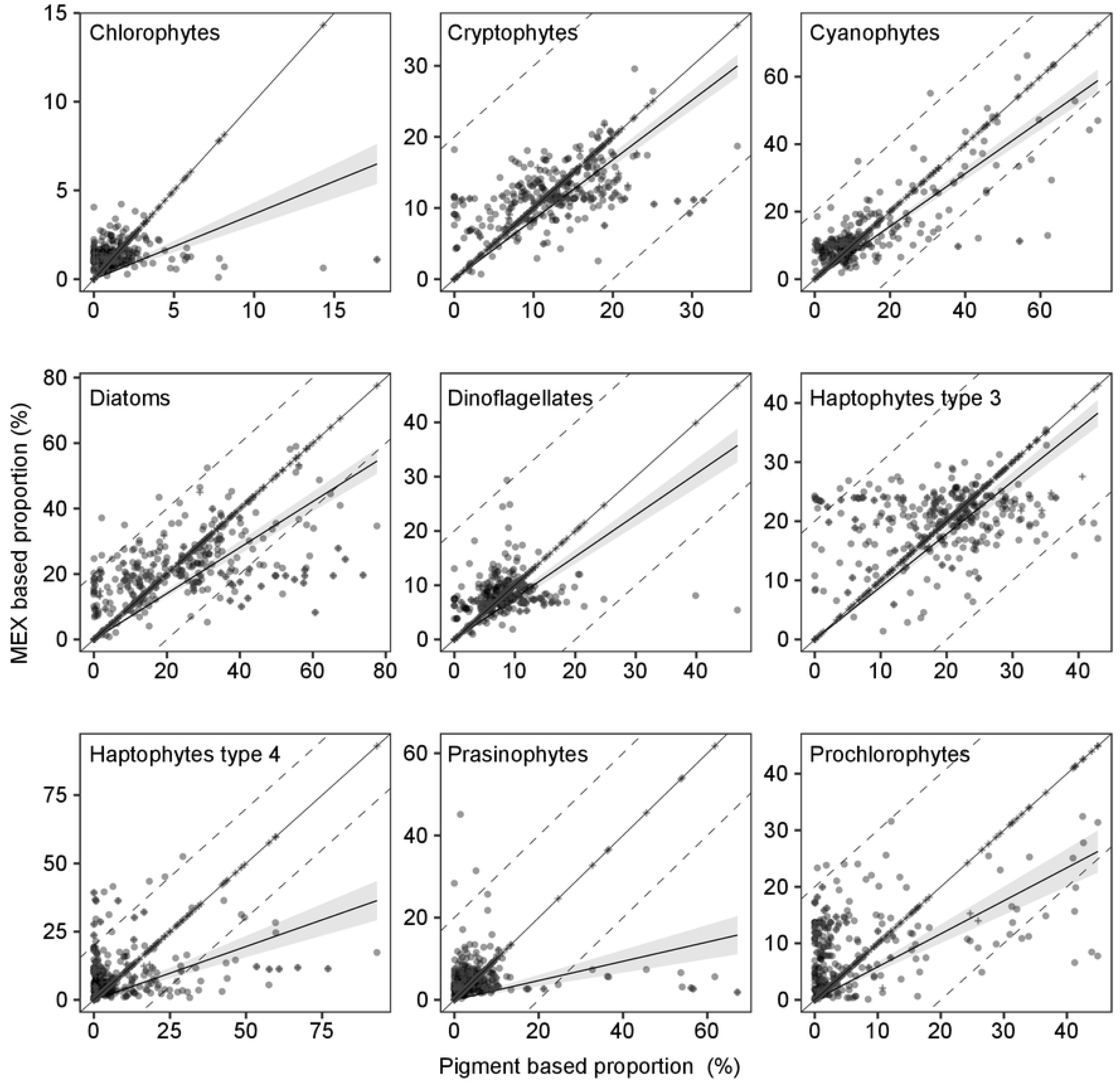
Comparisons of MEX (Multi-Exciter) - and plant-pigment-based proportions of TChla (total chlorophyll a concentration). Comparisons were performed in nine chemotaxonomic phytoplankton groups (chlorophytes, cryptophytes, cyanophytes, diatoms, dinoflagellates, haptophyte types 3 and 4, prasinophytes, and prochlorophytes). The thin solid line, dashed lines, and the bold line with shadow denote the 1:1 line, 20% fraction range relative to the 1:1 line, and the linear regression line at 95% confident intervals (intercept: 0), respectively. Linear regression was used only for data calculated from the database excluding the same station.

Sensitivity analysis indicated that the proportion of every chemotaxomic group showed a significant positive relationship between MEX estimation and plant-pigment-based estimations when the intercept was fixed at 0 (*p* < 0.001, *r*^2^ > 0.145) (Fig 2). We observed weak relationships (*r*^2^ < 0.5) in chlorophytes, type 4 haptophytes, and prochlorophytes. When the intercept was not fixed at 0, we observed no significant positive relationships between MEX estimation and plant-pigment-based estimations in chlorophytes and prasinophytes. In chlorophytes, the MEX-estimated proportion was always ≤5%, whereas the plant-pigment-based proportion sometimes recorded a proportion ≥5%. For prasinophytes, the proportion was largely <20%, and a significant positive relationship was observed when the data were limited, as both MEX-estimated and plant-pigment-based proportions were <20%.

When focused on chlorophyll *a* concentration-based abundance, we observed significant positive relationships between MEX and plant-pigment analyses (*p* < 0.001), except for type 4 haptophytes and prochlorophytes (Fig 3). For plant-pigment analysis, both chlorophyll-based type 4 haptophyte and prochlorophyte abundances were <0.2 µg L^−1^, whereas MEX estimations sometimes showed abundances >0.25 µg L^−1^. In the other chemotaxomic groups, we found overestimated results in MEX estimations as compared with plant-pigment analysis. However, the coefficients of linear regressions indicated that the MEX-estimated chlorophyll *a* concentration was underestimated relative to that from plant-pigment analysis, except for cyanophytes and type 3 haptophytes.

**Fig 3.**
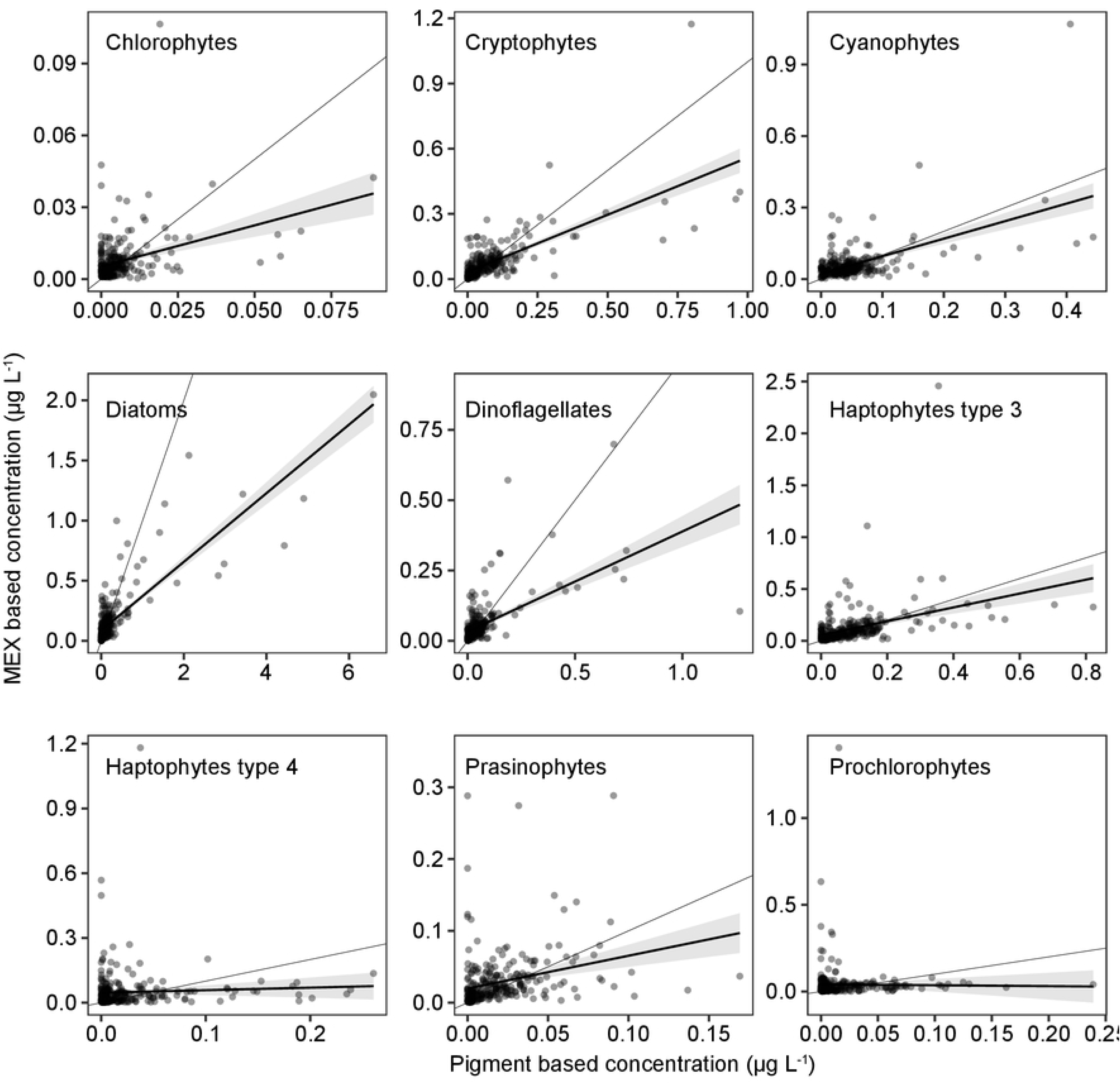
Comparisons of plant-pigment-based and MEX estimations of chlorophyll *a* concentration. Comparisons in nine chemotaxonomy phytoplankton groups (chlorophytes, cryptophytes, cyanophytes, diatoms, dinoflagellates, haptophyte types 3 and 4, prasinophytes, and prochlorophytes). The thin solid line, and the bold line with shadow denote the 1:1 line, and the linear regression line at 95% confident intervals (intercept: 0), respectively. Linear regression was used only for data calculated from the database excluding the same station.

Sensitivity analysis results suggested that the proportions and abundances of diatoms, dinoflagellates, cryptophytes, and cyanophytes were well estimated by the MEXs, whereas chlorophytes, prasinophytes, type 4 haptophytes, and prochlorophytes were not. Therefore, we summarized the proportion and abundance of prochlorophytes along with those of cyanophytes (named as cyanobacteria), and chlorophytes, prasinophytes, and type 4 haptophytes were summarized along with those of type 3 haptophytes (named as eukaryotes), although the abundance and proportion of type 3 haptophytes type were well estimated by the MEX. The MEX-estimated eukaryotes and cyanobacteria concentrations and abundances were positively correlated with those estimated by plant-pigment analyses according to sensitivity analysis (Fig 4).

**Fig 4.**
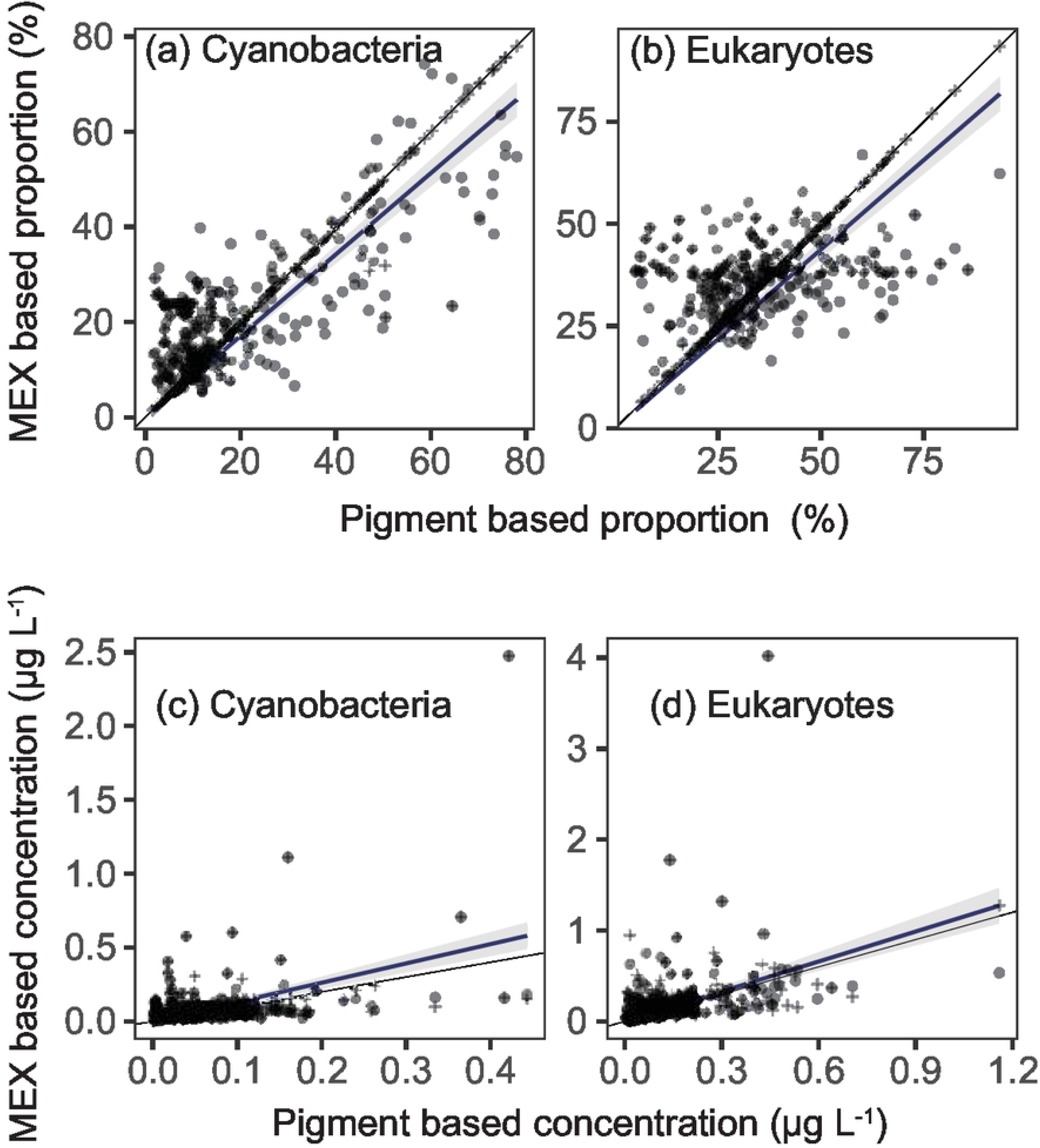
Comparisons of MEX (Multi-Exciter)- and plant-pigment-based proportions and chlorophyll *a* concentrations. Comparisons included MEX- and plant-pigment-based proportions of (a) cyanobacteria (sum of cyanophytes and prochlorophytes) and (b) eukaryotes (sum of chlorophytes, prasinophytes, and haptophyte types 3 and 4) and MEX- and plant-pigment-based chlorophyll *a* concentrations in (c) cyanobacteria and (d) eukaryotes. Crosses and circles denote MEX-based proportions calculated from the database containing and excluding the same stations, respectively. The thin solid line and the bold line with shadow denote the 1:1 line and the linear regression line at 95% confident intervals (intercept: 0), respectively. Linear regression was used only for data calculated from the database excluding the same station.

### Spatial and seasonal plant-pigment-based phytoplankton assemblages

The seasonal, spatial, and vertical distributions of plant-pigment-based chemotaxonomies of phytoplankton assemblages are shown in Fig 5. The vertical profiles were averaged every 25 m at depths of 0 m to 12.5 m and 12.5 m to 37.5 m. The seasonal variations showed that the TChla was highest during spring in all seasonally-observed areas. During spring, the diatom contribution was ∼50%, and the contribution of cyanobacteria (prochlorophytes + cyanophytes) was ≤10% in all areas (Fig 5). During the other seasons, there were significant spatial variations in the phytoplankton assemblages. At first, TChlas were higher in the Oyashio area, with the same levels observed in the JS and Kuroshio area, except in the winter, during which the TChla was higher in the Kuroshio area than the JS (Fig 5). The plankton assemblages showed that diatoms were dominant, with their contribution >50% at a depth of 50 m (at the base of the euphotic layer) in Okhotsk. Additionally, cyanobacteria (prochlorophytes + cyanophytes) significantly contributed (>50%) to the surface layers (≥30 m) during summer in the Kuroshio.

**Fig 5.**
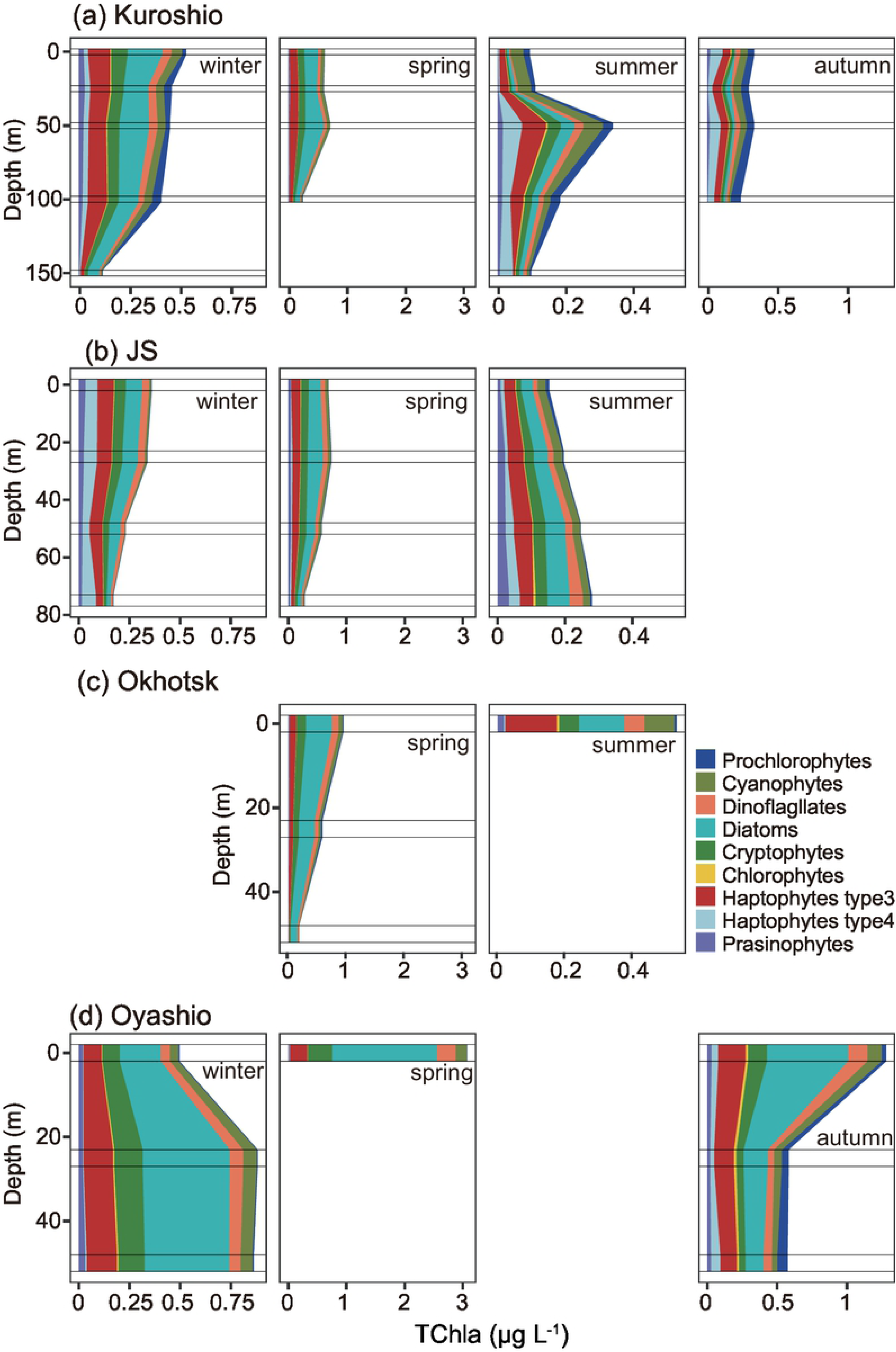
Seasonal and spatial variations in the vertical distributions of plant-pigment-based chemotaxonomic abundances. Concentrations were averaged every 25 m in depth.

PERMANOVA indicated significant differences in the chemotaxomic phytoplankton groups according to temperature and depth (*p* < 0.01, *R*^2^ = 0.13 and 0.02; respectively). In addition to temperature and depth, the nine light-source-excited fluorescence measurements explained the significant differences observed in the chemotaxomic phytoplankton groups (*p* < 0.05). The detection coefficient of every light source differed among the equations, and the sum of the detection coefficients was 0.37 (residuals: *R*^2^ = 0.63).

### Vertical distributions of MEX-estimated phytoplankton assemblages

The MEX observations showed significant seasonal and horizontal (area) differences, as well as plant-pigment-based differences (Fig 6). In the Kuroshio area, the contribution of cyanobacteria (prochlorophytes + cyanophytes) was high in the surface layers during summer and autumn, whereas it was low in winter and spring (Fig 6). The contributions of diatoms and cryptophytes varied seasonally, with their contributions high during spring and winter and low during summer. Contributions of eukaryotes (chlorophytes + prasinophytes + haptophytes) and dinoflagellates were relatively constant as compared with the other three groups. The contribution of dinoflagellates was <10%, whereas that of eukaryotes was usually >40%, except in the surface layer during summer (Fig 6).

**Fig 6.**
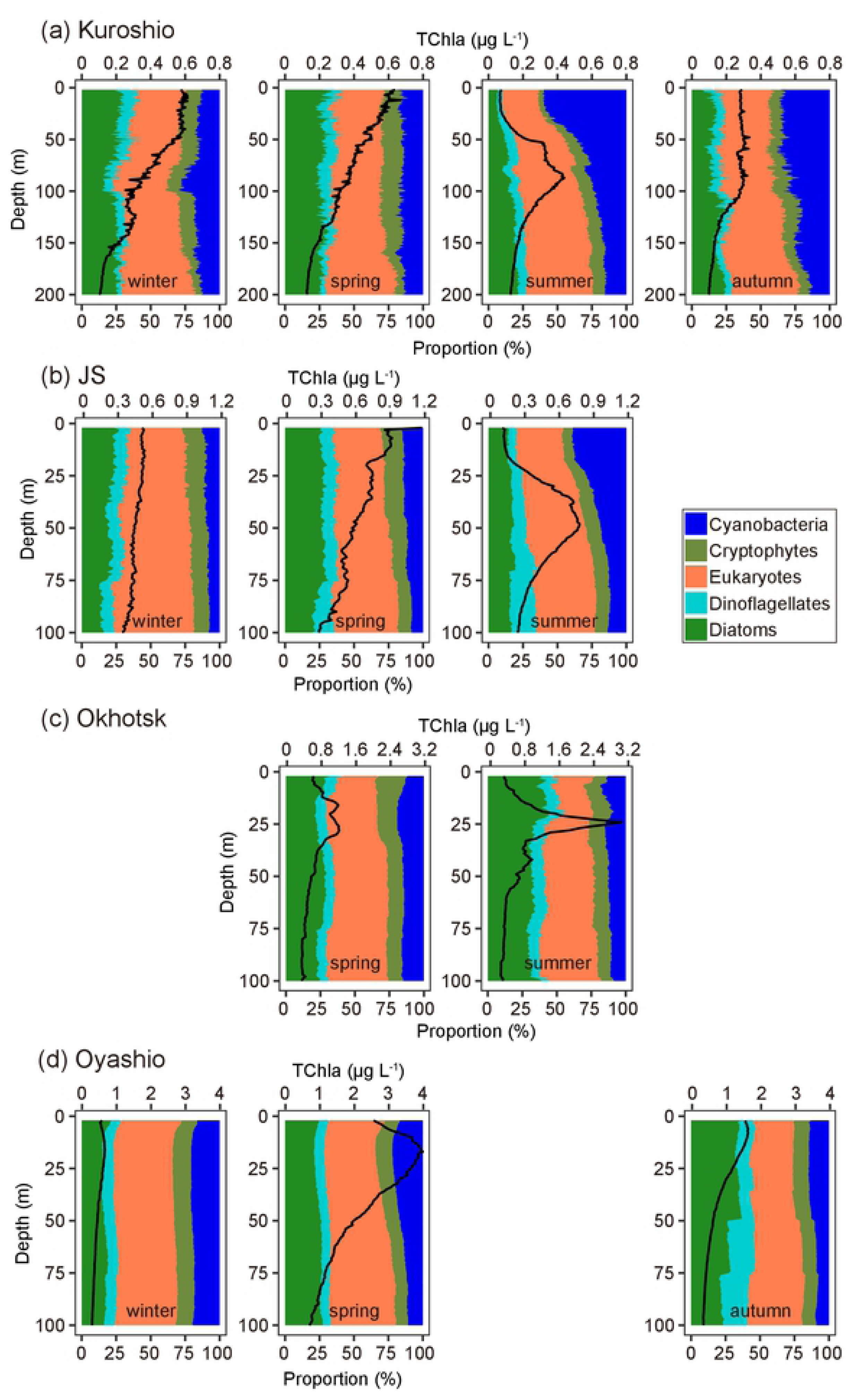
Vertical distributions of MEX (Multi-Exciter)-based mean chemotaxonomic proportions according to season. Distributions for (a) Kuroshio, (b) Sea of Japan (JS), (c) Okhotsk Sea, and (d) Oyashio, including the Oyashio–Kuroshio transition zone (Oyashio). The solid black line represents the MEX-based mean total chlorophyll a concentration (TChla).

In the JS, the phytoplankton assemblages showed similar seasonality to those of the Kuroshio (Fig 6). The diatom contributions were high during spring (April), and the contribution of cyanobacteria was high during summer in the surface layer. However, he contribution of cyanobacteria showed annual variation, as its contribution was >30% in 2019 and <25% in 2020 (data not shown). The eukaryotes were the dominant group at all seasons and depths. Interestingly, the dinoflagellate contribution was the same (∼15%) as that of diatoms below the subsurface chlorophyll maximum during summer (Fig 6).

In the Oyashio area, seasonal variations differed from those of the JS and Kuroshio, with diatoms dominant during winter and summer and eukaryotes at the same level as the diatoms in autumn (Fig 6). Additionally, the contributions of cryptophytes, cyanobacteria, and dinoflagellates were similar vertically and seasonally, except for dinoflagellates during autumn, with the contribution of dinoflagellates elevated at the subsurface. In addition to seasonal variations, we observed significant horizontal variations in the Oyashio area, particularly during autumn. In the coastal area, diatoms were dominant, whereas in the offshore area, cyanobacteria were dominant on the surface.

In Okhotsk, the plankton assemblages were similar in June (spring) and September (summer), and diatoms were dominant, followed by eukaryotes. During spring, the contribution of cryptophytes was high near the surface (Fig 6).

### Water-column-integrated phytoplankton abundance

The integrated chemtaxonomic TChla in the water column (2–200 m in the Kuroshio; and 0–100 m in the other three areas) indicated that eukaryotes were most abundant in the Kuroshio and JS areas throughout the year, whereas diatoms were abundant in the Okhotsk and Oyashio (Fig 7). The water-column-integrated diatom abundance was highest during spring, followed by winter in the Kuroshio, JS, and Oyashio (Fig 7). In the JS and Kuroshio areas, diatom abundance differed significantly (ANOVA with Tukey’s test; *p* < 0.05) during spring and summer, but these differences were not significant during winter and spring (*p* > 0.05). No significant difference was observed in diatom abundance between spring and summer in the Okhotsk (*t* test; *p* > 0.05).

**Fig 7.**
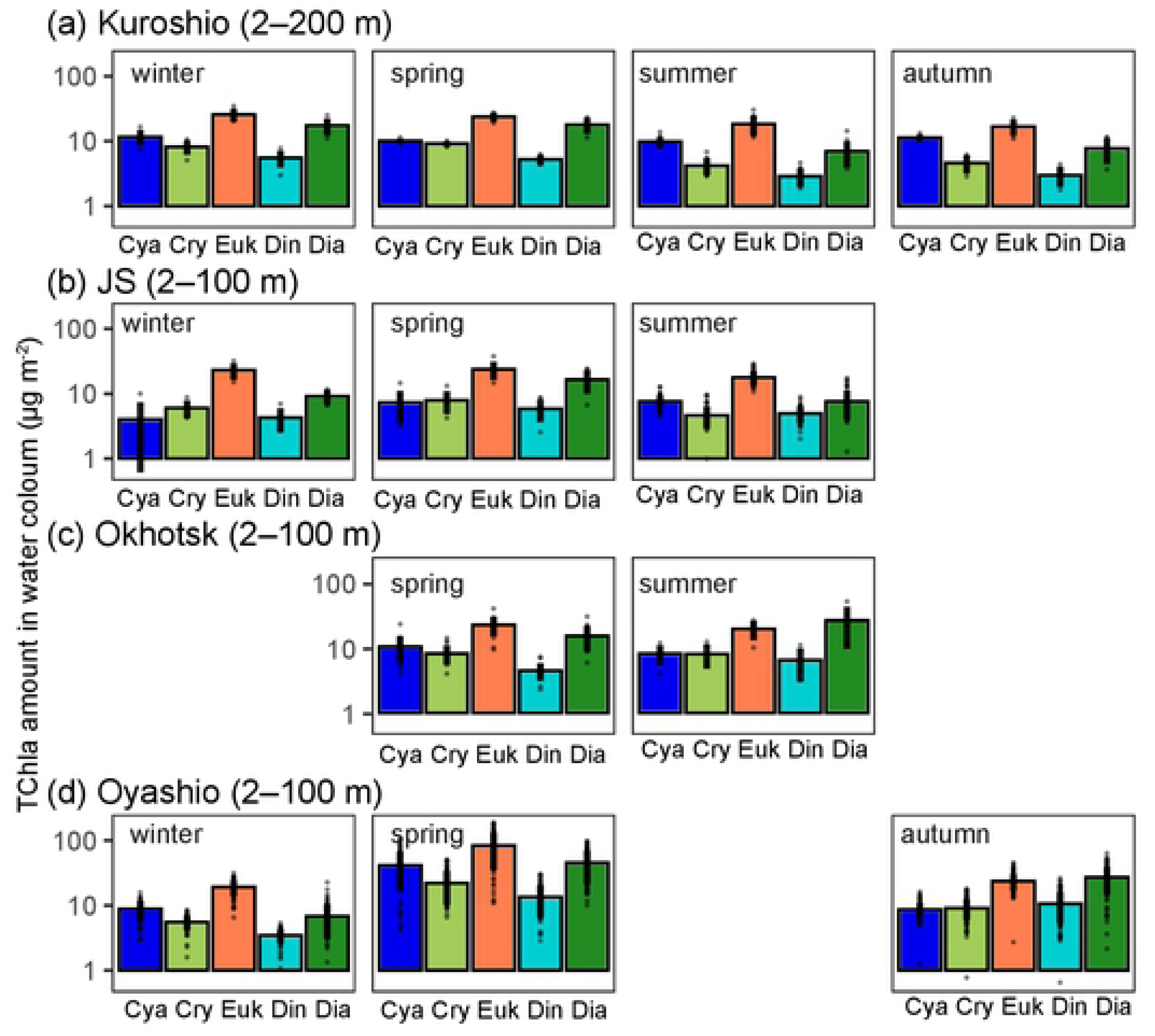
Seasonal and spatial variations in water-column-integrated abundances and MEX (Multi-Exicter)-based chemotaxonomic abundances. Water-column-integrated abundances (2-200 m in the Kuroshio area; and 2–100 m in the other three areas). The Y-axis represents log-scale values. Cya: cyanobacteria; Cry: cryptophyes; Euk: eukaryotes; Din: dinoflagellates; Dia: diatoms.

Water-column-integrated eukaryotes showed a similar seasonal variation as that of diatoms in the Kuroshio area, with winter the highest and significantly higher than those during summer and autumn (ANOVA with Tukey’s test; *p* < 0.05), although no significant difference was observed with that of spring (Fig 7). In the JS, water-column-integrated eukaryote abundance showed significant seasonal variation (ANOVA; *p* < 0.05), although a significant difference between seasons was not observed (ANOVA with Tukey’s test; *p* > 0.05). In the Oyashio area, water-column-integrated eukaryote abundance was at the same level during spring and autumn, and that in winter was significantly lower than that in spring and autumn (ANOVA with Tukey’s test; *p* < 0.05) (Fig 7).

Water-column-integrated cyanobacteria abundance did not show seasonal variations in the Kuroshio area (ANOVA; *p* > 0.05) (Fig 7). In the JS, cyanobacteria abundance during spring and summer was significantly higher than that in winter (ANOVA with Tukey’s test; *p* < 0.05). In the Oyashio area, cyanobacteria abundance was highest in spring, with significant differences observed between summer and winter (ANOVA with Tukey’s test; *p* < 0.05).

Seasonal vaticinations of water-column-integrated cryptophyte and dinoflagellate abundance were similar to that of diatoms in the Kuroshio area and the JS (Fig 7), with those in winter and spring significantly higher than those in summer and autumn (ANOVA with Tukey’s test; *p* < 0.05). In the Oyashio area, water-column-integrated cryptophyte abundance was significantly higher in the spring (ANOVA with Tukey’s test; *p* < 0.05) but did not differ significantly between autumn and winter (ANOVA with Tukey’s test; *p* > 0.05), although that of dinoflagellates was significantly higher in autumn relative to winter (ANOVA with Tukey’s test; *p* < 0.05).

### Comparison of surface and water-column-integrated abundance

We then compared the surface concentration and water-column mean concentrations by dividing the water-column mean values [98 m, except in the Kuroshio area (198 m)] by the surface values (at a mean depth of 2–5 m), followed by transformation into logarithmic values. This ratio (described as the column:surface TChla ratio) was 0 when the surface abundance was equal to that at a depth of 100-m (or 200 m for the Kuroshio area), and positive values indicated that the water-column-integrated amount was underestimated based on the surface values. We found that the column:surface TChla ratio differed according to season, with the ratio elevated during summer in areas observed during that season (Fig 8). Additionally, this ratio was relatively low for cyanobacteria but high in the other four chemotaxonomic groups during summer (Fig 8). During the other seasons, the differences among chemotaxonomic groups were unclear.

**Fig 8.**
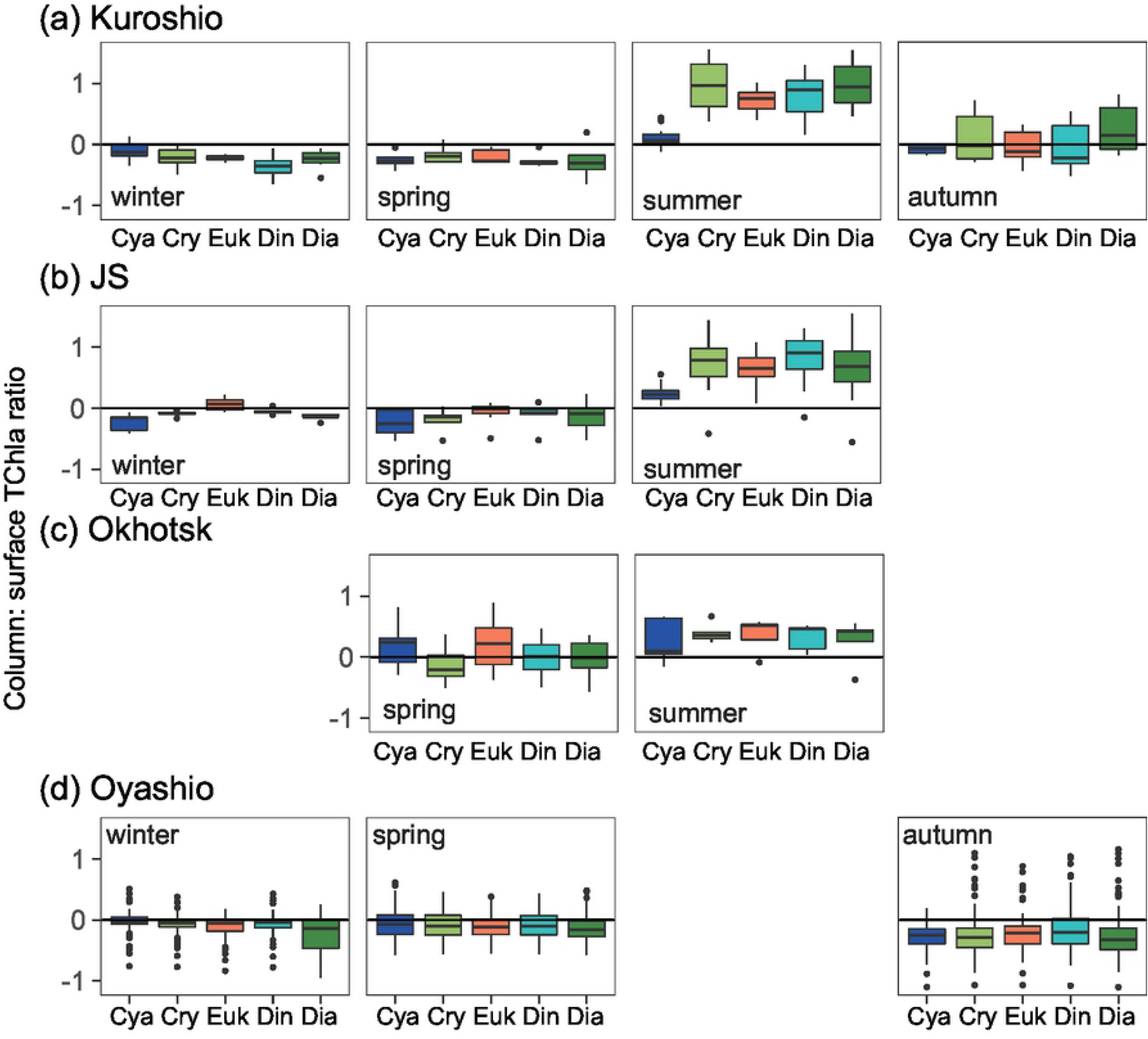
Seasonal and spatial variations in the water-column mean TChla:surface TChla ratio in all chemotaxomical groups. Water-column mean TChla measurements were made at a depth between 2 m and 100 m, except for the Kuroshio area (2–200 m). TChla: total chlorophyll *a* concentration; Cya: cyanobacteria; Cry: cryptophyes; Euk: eukaryotes; Din: dinoflagellates; Dia: diatoms.

## Discussion

In this study, we empirically estimated the chlorophyll-based abundances of five chemotaxonomic phytoplankton groups to a sensitivity greater than that previously reported [7, 8, 10]. Specifically, the method used in the present study distinguished the contributions of diatoms and dinoflagellates. Yentsch and Yentsch (21)] showed that the excitation spectra of diatoms and dinoflagellates are similar; however, Johnsen, Prézelin (22)] showed that the peridinin–chlorophyll *a*-binding protein complex demonstrated a high degree of fluorescence when excited at between 480 nm and 510 nm (peridinin is a marker pigment of dinoflagellates [1]). These results suggest that the contributions of diatoms and dinoflagellates were possibly detected by the MEX at an excitation wavelength of 505 nm using an LED, whereas there is no light source attributed to this wavelength in the Fluoroprobe.

The conversion methods applied in this study enabled evaluation of five chlorophyll-*a*-based chemotaxonomic groups; however, the results also demonstrated limitations of the MEX. MEX data, including pressure and temperature, only explained 37% of plant-pigment chemotaxonomic data based on PERMANOVA, suggesting that the MEX observations theoretically explained 37% of the variations in the phytoplankton assemblage. The results of PERMANOVA indicated that the quality of the estimation did not improve significantly when sensor depth was considered during the conversion processes. The empirical conversion method used here adequately converted the fluorescence to phytoplankton assemblages. Therefore, the expansion of the reference database is necessary to ensure improved evaluations, and we recommend that some plant-pigment samples be collected with MEX observations.

The shortcomings of the MEX observations were obvious, although our results showed significant spatial-seasonal variations in phytoplankton assemblages in the vicinity of Japan. The MEX results corresponded to area-specific features: the abundance of diatoms was high in the Oyashio area, particularly during spring [23, 24]. Additionally, the high contribution of cyanobacteria in the Kuroshio area during summer near the surface was the same as that in the more coastal and offshore areas under the influence of the Kuroshio area [18, 25]. Moreover, the high diatom and cryptophyte contributions in the Kuroshio area are similar to those reported previously [18, 19, 25].

The present study is the first to report phytoplankton assemblages for the eastern part of the JS. The abundance and contributions of diatoms were high in spring, which is consistent with the microscopic observations from the central part of the JS [26]. Additionally, we found that the contribution of cyanobacteria was high at the surface of the JS during summer, as well as in the Kuroshio area. Inoue et al. [27] noted that >60% of the surface water of the southwestern JS originated in the Kuroshio area based on the radium isotope ratio, and that macro-nutrients are depleted at the surface of both the JS and the Kuroshio area during summer [28, 29]. We considered that these environmental factors induced cyanobacteria-rich surface waters in the eastern JS. However, the importance of eukaryotes, which comprise pico- and nano-sized eukaryotic phytoplankton, was not indicated; therefore, the role of the eukaryotic phytoplankton community in the JS should be studied in the future.

MEX allows calculation of the water-column-integrated abundance of every group. In previous studies, discrete samples were essential for estimating phytoplankton assemblages; therefore, the water-column-integrated abundance was hardly ever calculated. However, Latasa et al. [30] found that phytoplankton contribution differs at a fine vertical scale. Therefore, interpolation of the discrete observations do not always reflect the water-column-integrated values. Bracher et al. [31] first estimated the water-column-integrated abundance using hyperspectral underwater-irradiance radiometer data in the Atlantic Ocean. These radiometers can only be applied during the day, which suggests an advantage for the MEX. The present study is the first report on water-column-integrated phytoplankton abundance at the chemotaxonomic group level in the Pacific Ocean. The column:surface TChla ratio indicated that the surface concentration did not always reflect the water-column-integrated abundance. In particular, we identified bias during the summer and involving cyanobacteria abundance, because the contribution of maximum subsurface chlorophyll was high during summer, and the cyanobacteria contribution was high above the subsurface chlorophyll maximum in the well-stratified water column. These observations were not conducted equally among areas; therefore, we did not evaluate spatial differences. Hirata et al. [4] reported surface phytoplankton abundance at the chemotaxonomic group level based on space-borne ocean-color sensors. In the present study, we showed that estimation of the water-column surface phytoplankton abundance (i.e., global-scale abundance at the chemotaxonomic group level) remains difficult. However, further observations using the MEX might enable estimations with higher accuracy.

The spatial-seasonal variations observed in this study were largely similar to the phytoplankton-community structure previously reported from discrete samples [18, 19, 23–25]. However, we obtained novel insights from the MEX observations, including pico- and nano-eukaryotic phytoplankton contributions in the JS across seasons. More detailed analyses of datasets with environmental parameters can allow a deeper understanding of primary productive processes in the ocean. The MEX is light and can be attached to autonomous platforms, which could enable estimation of the global-scale abundance at the chemotaxonomic group levels in near future. Additionally, further developments in the MEX might allow acquisition of primary production rates at the chemotaxonomic group level. Kazama at al. [32] reported the gross primary productivity at the chemotaxonomic group level in a freshwater lake using a machine with similar capabilities, although that method used the fluorescence yield of the photosystem II complex under background light.

## Conclusion

In this study, we described a method allowing reliable estimation of phytoplankton-community structure using the MEX and showed spatial-seasonal variations of vertical profiles and water-column-integrated values. Notably, the reference data were limited to those from the vicinity of Japan; therefore, the results suggest the effectiveness of this method exclusively in coastal and offshore waters around Japan. However, this approach represents a promising method for potential worldwide after the expansion of the reference datasets. Moreover, although validation is required, this approach is not limited to the MEX but can also be applied to other submarine spectrofluorometers, such as Fluoroprobe.

## Acknowledgments

We thank the captains, crews, and scientists of RVs *Soyo-Maru*, *Hokko-Maru*, *Wakataka-Maru,* and TV *Tenyo-Maru* of the Japan Fisheries and Research Education Agency and *Dai-5 Kaiyo-Maru* of Kaiyo Engineering Co., Ltd. for their invaluable support with sampling.

